# Polybasic patches in both C2 domains of Synaptotagmin-1 are required for evoked neurotransmitter release

**DOI:** 10.1101/2021.07.05.451149

**Authors:** Zhenyong Wu, Lu Ma, Nicholas A. Courtney, Jie Zhu, Yongli Zhang, Edwin R. Chapman, Erdem Karatekin

**Author notes:** see 4 (Zhenyong Wu). Institute of Physics, Chinese Academy of Sciences, Beijing, China (Lu Ma). Oklahoma Medical Research Foundation, 825 NE 13th Street, Oklahoma City, Oklahoma 73104. These authors contributed equally to this work (Zhenyong Wu, Lu Ma).

## Abstract

Synaptotagmin-1 (Syt1) is a vesicular calcium sensor required for synchronous neurotransmitter release. It is composed of a single-pass transmembrane domain linked to two tandem C2 domains (C2A and C2B) that bind calcium, acidic lipids, and SNARE proteins that drive fusion of the synaptic vesicle with the plasma membrane. Despite its essential role, how Syt1 couples calcium entry to synchronous release is not well understood. Calcium binding to C2B, but not to C2A, is critical for synchronous release and C2B additionally binds the SNARE complex. The C2A domain is also required for Syt1 function, but it is not clear why. Here we asked what critical feature of C2A may be responsible for its functional role, and compared this to the analogous feature in C2B. We focused on highly conserved poly-lysine patches located on the sides of C2A (K189-192) and C2B (K324-327). We tested effects of charge-neutralization mutations in either region (Syt1^K189-192A^ and Syt1^K326-327A^) side-by-side to determine their relative contributions to Syt1 function in cultured cortical mouse neurons and in single-molecule experiments. Combining electrophysiological recordings and optical tweezers measurements to probe dynamic single C2 domain-membrane interactions, we show that both C2A and C2B polybasic patches contribute to membrane binding, and both are required for evoked release. The readily releasable vesicle pool or spontaneous release were not affected, so both patches are specifically required for synchronization of release. We suggest these patches contribute to cooperative binding to membranes, increasing the overall affinity of Syt1 for negatively charged membranes and facilitating evoked release.

**Significance Statement:** Synaptotagmin-1 is a vesicular calcium sensor required for synchronous neurotransmitter release. Its tandem cytosolic C2 domains (C2A and C2B) bind calcium, acidic lipids, and SNARE proteins that drive fusion of the synaptic vesicle with the plasma membrane. How calcium-binding to Synaptotagmin-1 leads to release and the relative contributions of the two C2 domains is not clear: unlike C2B, calcium-binding to C2A is not critical for evoked release, yet both domains are needed for Syt1 function. Combining electrophysiological recordings from cultured neurons and optical tweezers measurements that probe single C2 domain-membrane interactions, we show that conserved polybasic regions in both domains contribute to membrane binding cooperatively, and both are required for evoked release, likely by increasing the overall affinity of Synaptotagmin-1 for negatively charged membranes.

## INTRODUCTION

Synaptotagmin-1 (Syt1) is a major neuronal calcium sensor for synchronous neurotransmitter release^1-5^. In mice, knock-out (KO) of *syt1* is lethal at birth, with nearly complete elimination of synchronous release from cultured hippocampal neurons collected from *syt1*^*-/-*^ newborns^4^. In humans, mutations in Syt1 result in severe neurodevelopmental disorders^6^. Syt1 is a synaptic vesicle protein comprising a short N-terminal luminal sequence, a single-pass transmembrane domain, and a cytoplasmic linker region followed by two tandem C2 domains (C2A and C2B) at the C-terminus that bind calcium, acidic lipids, and SNARE proteins^5^. Despite its critical role in neurotransmitter release, how Syt1 couples calcium entry to synchronous membrane fusion is not well understood^5,7-10^. In particular, the relative contributions of the two tandem C2 domains of Syt1 to the overall function of the protein are not clear. Here we show that conserved polybasic patches in both C2 domains are required for evoked release and efficient membrane binding.

The C2 domains of Syt1 are 8-stranded β-sandwich structures with two protruding loops that form the Ca^2+^ binding pockets, with 3 and 2 Ca^2+^ ions binding to C2A and C2B, respectively, *via* a series of highly conserved aspartates^5^. Calcium binding leads hydrophobic residues at the tips of the calcium-binding loops to bury into the membrane for both C2A and C2B, with preference for bilayers containing phosphatidylserine (PS) or phosphatidylinositol 4,5-bisphosphate (PI(4,5)P_2_) for the former and latter, respectively^11-15^. Disruption of calcium binding to C2B, but not to C2A, eliminates synchronous neurotransmitter release, suggesting calcium binding to C2B, but not to C2A, is essential for triggering synchronous release^3,16-20^. C2 domains also interact with SNARE proteins in both calcium-dependent and independent manner, but there is still some debate as to which interactions are most relevant physiologically^21-25^. Early biochemical^26^ and recent structural studies suggest only the C2B domain binds the neuronal SNARE complex, that this binding mode is calcium-independent^25,27^, and maintained in the presence of membranes^28,29^.

Both the C2A and C2B domains have a highly conserved poly-lysine patch located on the side: K189-192 for the former and K324-237 for the latter (Figure 1A, B). The role of the polybasic patch in C2B has been studied in the past, with conflicting results. Reist and colleagues^30,31^ showed that replacing three of the C2B domain polybasic region lysines with glutamines (*syt1*^*K379,380,384Q*^) in *Drosophila* 3rd instar larvae resulted in a ∼40% decrease in evoked release and a doubling of spontaneous release at the neuromuscular junction (NMJ), effects attributed to vesicle priming defects in the mutant. Using autaptic cultures of *syt1* knock-out neonatal mice hippocampal neurons expressing wild-type or mutant Syt1, Li et al.^32^ found that partial neutralization of the C2B domain polybasic region of Syt1 (K326A, K327A, the “KAKA substitution”), similarly resulted in a 50% decrease in evoked release, but with little effect on spontaneous release or the readily releasable pool (RRP) size. Using a similar preparation, Borden et al.^33^ also reported mild effects for the K326A, K327A substitution: a ∼50% reduction in the fraction of vesicles released per stimulus and an unchanged RRP size. Both studies also found a reduction in the normalized EPSC amplitude as a function of extracellular calcium concentration. By contrast, rescue of *syt*^*-/-*^ neonatal autaptic hippocampal neuronal cultures with a Syt1 K325A,K327A mutant resulted in a much more severe, 5-6-fold reduction in synchronous release and a ∼60% reduction in the RRP size^34^. Chang et al. also found this mutation disrupts tight attachment of synaptic vesicles to the active zone plasma membrane^34^. It is not clear if the discrepancies among the reported results are due to differences in the species used, the preparation of the cultures, and/or the stimulation methods used.

**Figure 1.**
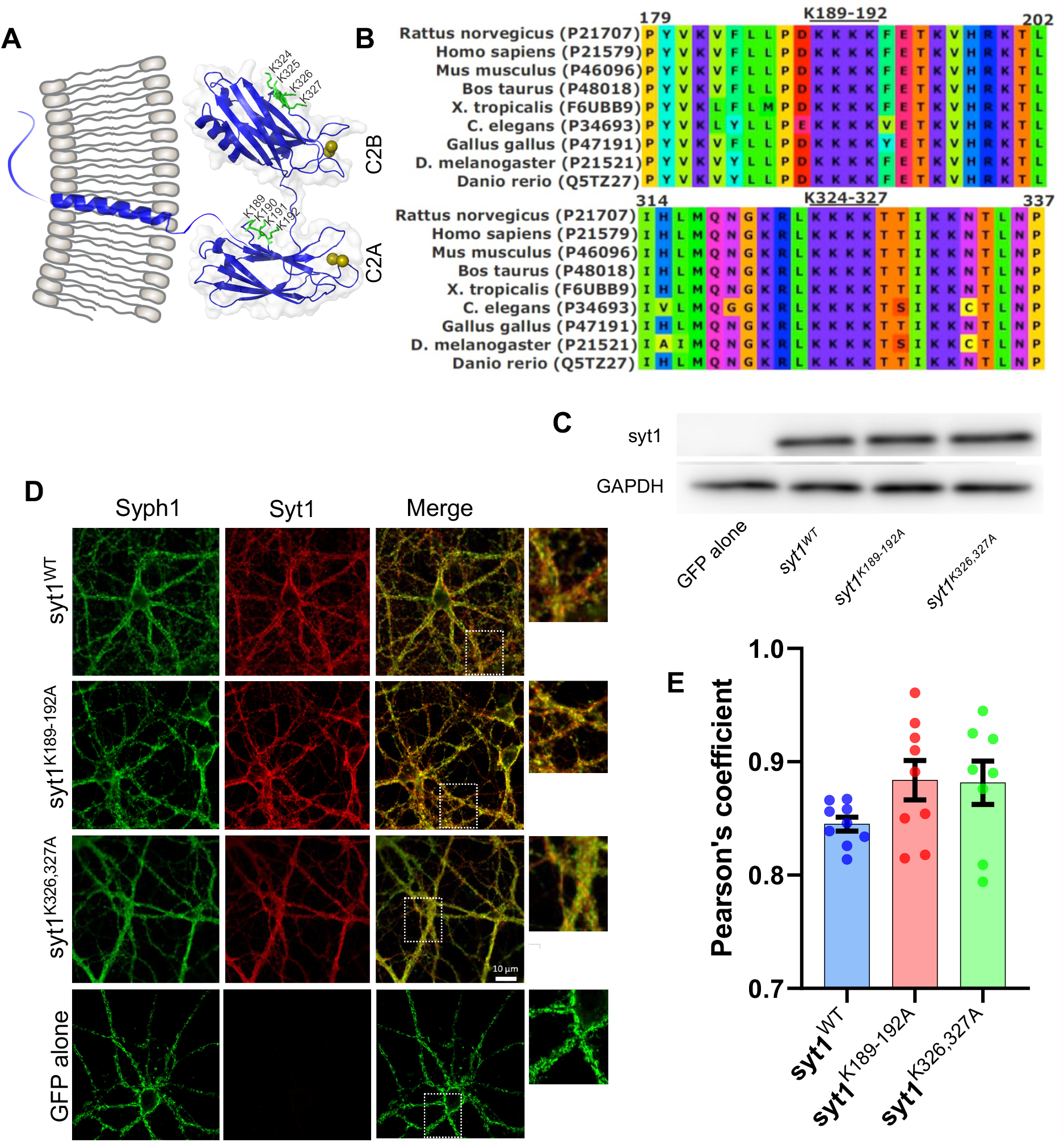
Expression and targeting of Syt1 polybasic patch mutants are similar to those of wild-type Syt1. **A**. Schematic of the structure of Syt1, with the polybasic patches marked with ball-and-stick representation in green. The numbering refers to the mouse sequence. Calcium ions are depicted as orange spheres. The C2A and C2B domains are rendered from PDB entry 5CCG using PyMOL^25^, while the rest of the molecule is schematically drawn using CorelDRAW. **B**. Multiple alignment of synaptotagmin-1 protein sequences from various species as indicated, using ClustalW^74^. The uniprot access codes are shown in parentheses (https://www.uniprot.org/). **C**. Western blot analysis of the expression of wild-type or mutant Syt1 transgenes in *syt1*^*-/-*^ mouse neonatal cortical cultures. A representative result from 3 separate experiments is shown. All *syt1* constructs were expressed at similar levels. **D**. Exogenously expressed wild-type or mutant Syt1 are correctly targeted. Immunofluorescence signals of Syt1 variants were compared with those of Synaptophysin-1 (Syph1), a synaptic vesicle marker. The boxed regions in the third column are shown expanded on the right. **E**. Quantification of colocalization of Syph1 and Syt1 immunofluorescence signals using the Pearson correlation coefficient (a value of 1 indicates perfect colocalization). There were no significant differences among the groups (one-way ANOVA, followed by Dunnett’s test to compare mutants against WT Syt1. WT vs. Syt1^K189-192A^: p=0.14, WT vs Syt1^K326327A^: p=0.18).

The C2A domain has received less attention since the finding that disrupting Ca^2+^ binding to C2A results in less severe phenotypes^16,18,19^ than a similar disruption in the C2B domain^17,20^. In addition, biochemical studies reported that the C2B domain, but not the C2A domain, binds phosphoinositides in a calcium-independent manner through its poly-lysine patch^11,14,15^. However, recent work suggests Ca^2+^ binding to C2A^35,36^, and the ensuing insertion of the hydrophobic residues at the tips of the C2A loops^37^ are required for evoked release, at least at the *Drosophila* larvae NMJ, consistent with biochemical experiments showing robust Ca^2+^-dependent membrane binding and penetration of the C2A domain to negatively charged membranes^5,13,28^. In addition, domain deletion and swapping experiments suggested that the C2A domain is essential for Syt1 function in *Drosophila*^38^ and more recently in mice^39^.

Despite these recent results showing the importance of the C2A domain, the function of the poly-lysine patch in C2A remains unclear. Neutralization of the C2A polybasic patch in *Drosophila* larvae NMJ resulted in a 2.4-fold increase in spontaneous release frequency but left evoked release intact^40^. By contrast, injection of a small peptide including the polybasic sequence into the squid giant nerve terminals inhibited transmitter release when the nerve was stimulated, suggesting this region to be important for evoked release^41^. Similarly, injection of the entire C2A domain into PC12 cells resulted in inhibition of Ca^2+^-triggered release, an effect attributed to the poly-lysine motif through alanine-scanning mutagenesis^42^.

In summary, the reported effects of mutations of the conserved polybasic patches in the two C2 domains of Syt1 are conflicting. We decided to test effects of charge-neutralization mutations in the two poly-basic domains side-by-side to determine their relative contributions to Syt1 function in cultured cortical mouse neurons and in biophysical single-molecule experiments. Combining electrophysiological recordings and single-molecule optical tweezers (OT) measurements to probe dynamic C2 domain-membrane interactions, we show that both C2A and C2B polybasic patches contribute to membrane binding, and both are required for evoked release. There is no measurable effect on the size of the readily releasable pool (RRP) of vesicles or spontaneous release, so both patches are specifically required for the synchronization of release. We suggest these patches contribute to cooperative binding to membranes, increasing the overall affinity of C2AB for negatively charged membranes and facilitating evoked release.

## RESULTS

### Synaptotagmin-1 polybasic patch mutants are expressed at wild-type levels and trafficked correctly

To assess the roles of the polybasic patches in its two C2 domains, we expressed Syt1 with mutations designed to neutralize the poly-lysine patch in the C2A (K189-192A) domain or the C2B (K326A,K327A) domain, designated as Syt1^K189-192A^ or Syt1^K326-327A^ respectively, in cultured cortical neurons from syt1^-/-^ mice^4^. We expressed wild-type Syt1 as a positive control or green fluorescent protein (GFP) alone as a negative control. Transgenes expressing wild-type or mutant Syt1 resulted in similar protein expression levels as assessed by Western blot analysis, whereas Syt1 was undetectable for the negative control (Figure 1C). As shown in Figure 1D,E, all three Syt1 variants were correctly trafficked to synaptic vesicles, as their immunofluorescence signals had a high correlation with those of Synaptophysin-1 (Syph1), a synaptic vesicle marker^39^.

### Both Synaptotagmin-1 C2A and C2B domain polybasic patches are required for evoked release

Next, we measured evoked release from cultured cortical neurons from Syt-1 KO mice expressing the transgenes using field stimulation (Figure 2A). Evoked release was nearly abolished in KO neurons expressing GFP alone (Figure 2B), consistent with previous reports^4^. Expression of wild-type Syt1 restored evoked release. By contrast, evoked release was greatly diminished in neurons expressing Syt1^K326-327A^. Interestingly, release was also largely reduced in neurons expressing Syt1^K189-192A^.

**Figure 2.**
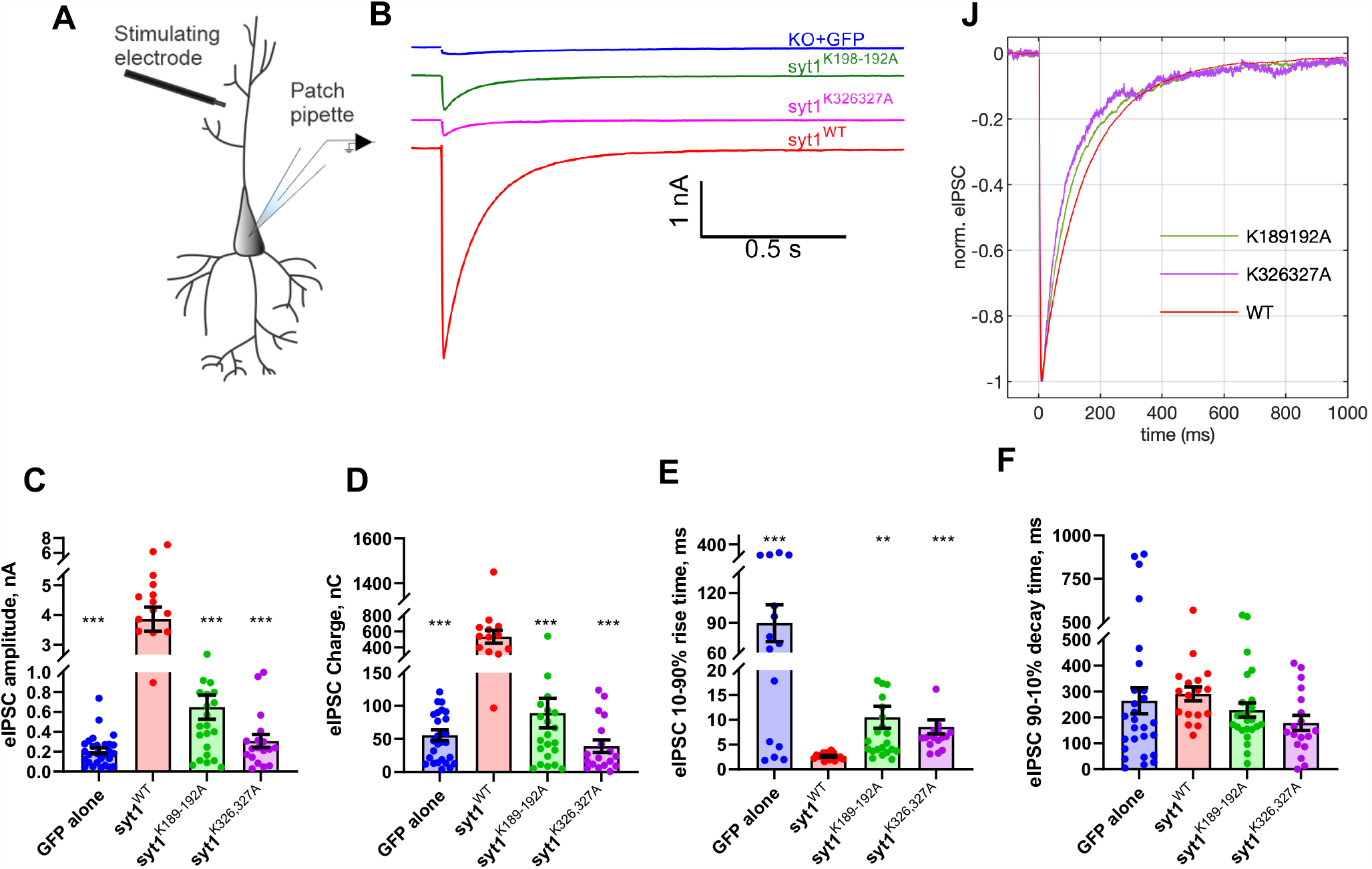
Syt1 C2A and C2B polybasic patch mutations dramatically reduce evoked release. **A**. Schematic of the recording configuration. **B**. Representative examples of inhibitory post-synaptic currents (IPSCs) recorded from cultured *syt1*^*-/-*^ cortical neurons expressing the indicated transgenes. Neurons expressing the Syt1 C2A (Syt1^K189-192A^) or C2B (Syt1^K326-327A^) polybasic patch mutations had greatly diminished responses compared to neurons expressing wild-type Syt1. Neurons lacking Syt1 expression had nearly all evoked release abolished. **C-D**. Quantification of evoked release parameters. **C**. eIPSC amplitudes were (mean ± S.E.M; in nA): GFP alone: 0.21 ± 0.03, WT: 3.9 ± 0.40, syt1^K189-192A^: 0.65 ± 0.12, syt1^K326,327A^: 0.31 ± 0.07. **D**. eIPSC charges (time integrals of the IPSC traces, mean ± S.E.M; in nC) were: GFP alone: 55.48 ± 8.12, WT: 531.5 ± 80.84, syt1^K189-192A^: 88.98 ± 22.38, syt1^K326,327A^: 38.97 ± 9.30. **E**. eIPSC rise times were (mean ± S.E.M; in ms): GFP alone: 55.48 ± 8.12, WT: 531.5 ± 80.84, syt1^K189-192A^: 10.21 ± 2.21, syt1^K326,327A^: 8.56 ± 1.41. **F**. eIPSC decay times were (mean ± S.E.M; ms): GFP alone: 263.8 ± 50.34, WT: 290.3 ± 26.80, syt1^K189-192A^: 228.1 ± 27.53, syt1^K326,327A^: 178.6 ± 29.19. (n=28, 17, 24, and 18 cells were tested for GFP alone, WT syt1, syt1^K189-192A^, and syt1^K326,327A^, respectively. Cells were prepared from N=5 *syt1*^*-/-*^ pups). Compared to neurons expressing Syt1^WT^, IPSC amplitudes (C) and charges (D) were lower by 6 and 12.6-fold for Syt1^K189-192A^ and Syt1^K326-327A^ neurons, respectively. The rise times of the Syt1^WT^, Syt1^K189-192A^, and Syt1^K326-327A^ constructs were shorter than for neurons lacking Syt1 (**E**), but the decay times were similar for all sets of neurons (**F**). There were no major kinetic differences among the averaged IPSCs normalized to the peak amplitude. For C-F, we used one-way ANOVA, followed by Dunnett’s test to compare mutants against WT Syt1. *, **, and *** indicate p<0.05, p<0.01, and p<0.001, respectively.

Quantification of evoked release parameters indicated the peak amplitude of the IPSCs and the charge transfer (integral of the IPSC traces) were significantly lower in the mutants compared with neurons expressing WT Syt1 (Figure 2C-D). There was no significant difference in the time to reach the IPSC peak or the decay time (Figure 2E-F). We normalized the amplitudes of averaged IPSCs to compare the kinetics of the evoked responses. Delays between stimulation and onset of release, and the release kinetics of normalized IPSCs were not significant for neurons expressing wild-type or mutant Syt1 (Figure 2J). These results indicate the poly-lysine patch in either C2 domain is critical for evoked release.

### Spontaneous release and the readily releasable pool of vesicles are not affected by Syt1 polybasic patch mutations

We next tested whether the polybasic patch mutations affected spontaneous release. Miniature IPSCs were recorded from resting *sty1*^*-/-*^ neurons expressing GFP alone, wild-type Syt1, or the polybasic patch mutants Syt1^K189-192A^ or Syt1^K326-327A^ (Figure 3). While the miniature IPSC frequency was elevated 2-3-fold in neurons lacking Syt1 as reported previously^43-46^, the frequency was similar in neurons expressing wild-type or mutant Syt1. Furthermore, there were no significant differences for miniIPSC amplitudes, rise times, or decay times among the experimental groups (Figure 3C-E). Thus, the polybasic patch mutations do not affect spontaneous release in this preparation.

**Figure 3.**
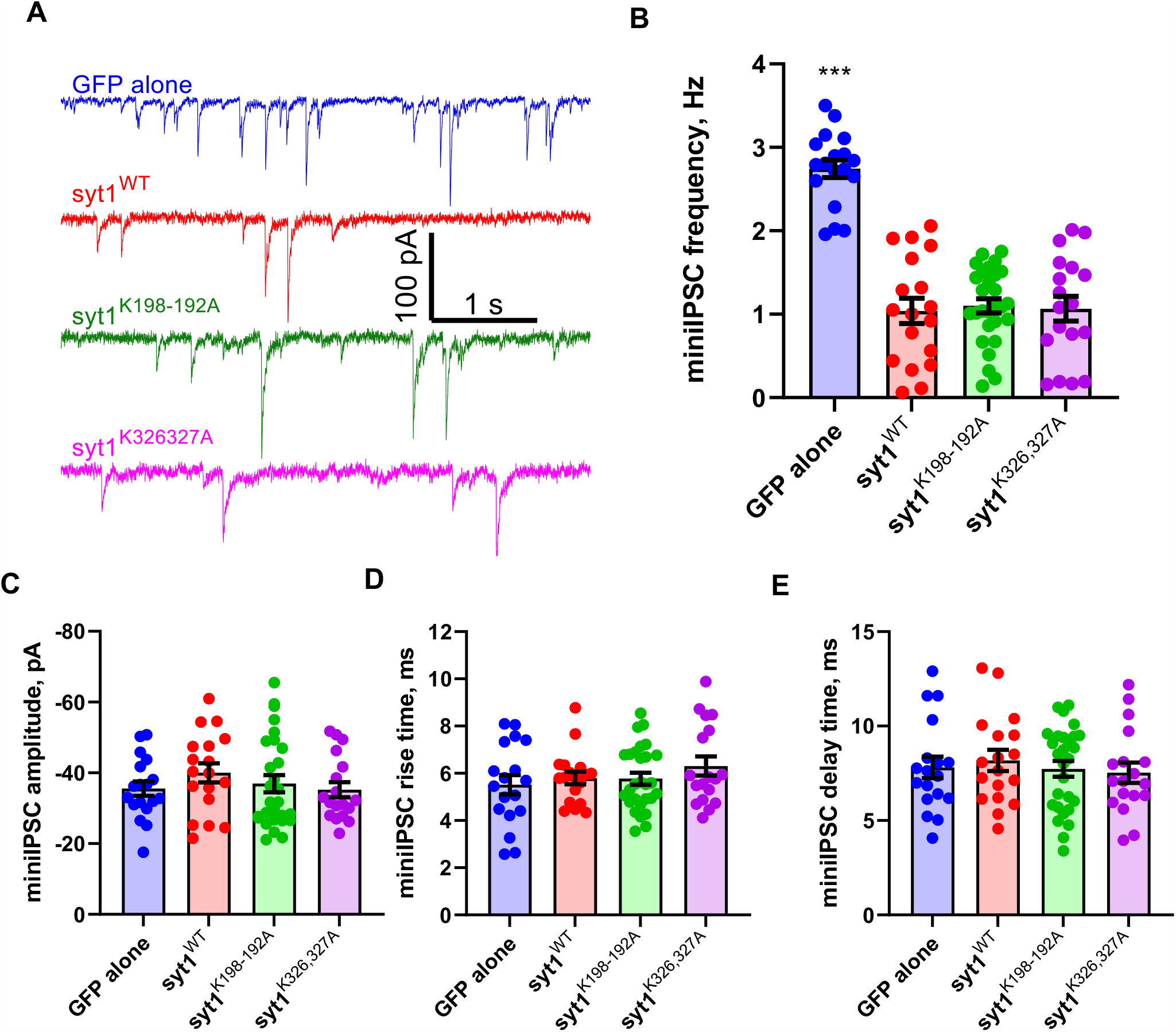
Syt1 C2A and C2B polybasic patch mutations do not affect spontaneous release. **A**. Representative current traces from voltage-clamped, resting cortical mouse *syt*^*-/-*^ neurons expressing the indicated transgenes. **B-D**. Quantification of miniature IPSC (miniIPSC) parameters from traces such as the ones shown in A. Apart from an increase in the miniIPSC frequency for neurons lacking Syt1 (**B**), there are no significant differences among the experimental groups for miniIPSC amplitude (**C**), rise time (**D**), or decay time (**E**). miniIPSC frequencies were (mean ± S.E.M): GFP alone: 2.74 ± 0.10, WT: 1.04 ± 0.15, syt1^K189-192A^: 1.1 ± 0.09, syt1^K326,327A^: 1.07 ± 0.15. miniIPSC charges were (mean ± S.E.M; pA) : GFP alone: 35.54 ± 2.04, WT: -40.00 ± 2.69, syt1^K189-192A^: -36.95 ± 2.42, syt1^K326,327A^: -35.22 ± 2.13. miniIPSC rise times were (mean ± S.E.M; ms): GFP alone: 5.16 ± 0.41, WT: 5.80 ± 0.27, syt1^K189-192A^: 5.77 ± 0.26, syt1^K326,327A^: 6.31 ± 0.40. miniIPSC decay times were (mean ± S.E.M; ms): GFP alone: 7.81 ± 0.57, WT: 8.183 ± 0.56, syt1^K189-192A^: 7.74 ± 0.42, syt1^K326,327A^: 7.536 ± 0.53. n=18, 18, 28, and 18 cells were tested for GPF alone, WT syt1, syt1^K189-192A^, and syt1^K326,327A^, respectively. Cells were prepared from N=3 *syt1*^*-/-*^ pups. For B-E, we used one-way ANOVA, followed by Dunnett’s test to compare mutants against WT Syt1. *, **, and *** indicate p<0.05, p<0.01, and p<0.001, respectively.

The reduction in evoked release we observed in neurons expressing the polybasic patch mutants Syt1^K189-192A^ or Syt1^K326-327A^ could be due to a defect in the size of the readily releasable pool (RRP) of synaptic vesicles. To test this possibility, we applied hypertonic sucrose to the four groups of neurons (Figure 4) and integrated the currents to estimate the total RRP size^47^. In neurons lacking Syt1, there was a 2-3-fold reduction in the RRP size, whereas the RRP sizes were similar for *syt1*^*-/-*^ neurons expressing wild-type Syt1 or the polybasic patch mutants. Thus, the polybasic patch mutations do not substantially affect RRP size measured by hypertonic sucrose application.

**Figure 4.**
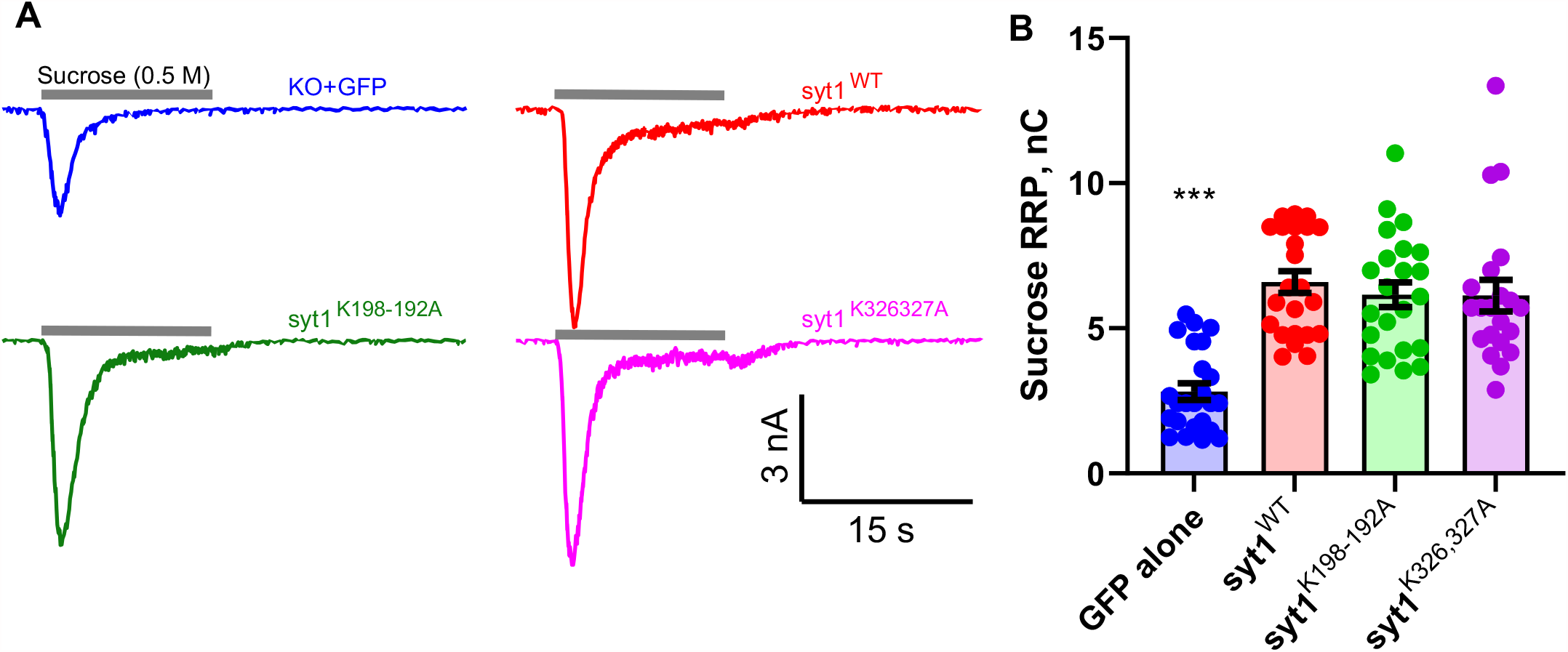
Polybasic patch mutations do not affect the readily releasable pool. **A**. Representative current traces elicited by application of a 0.5 M sucrose solution for 15 s, indicated by the gray bars above each trace, for syt1 KO neurons expressing the indicated transgenes. **B**. Integral of the hypertonic sucrose-induced currents to estimate the size of the readily releasable pool (RRP). The RRP size is indistinguishable for Syt1^WT^, Syt1^K189-192A^, and Syt1^K326-327A^, but is lower by about 2.4-fold for neurons lacking Syt1. Sucrose-induced total charges (RRP sizes) were (mean ± S.E.M; nC): GFP alone: 2.82 ± 0.28, WT: 6.59 ± 0.375, syt1^K189-192A^: 6.16 ± 0.43, syt1^K326,327A^: 6.13 ± 0.54. (n=24, 23, 23, and 21 cells were tested for GFP alone, syt1 WT, syt1^K189-192A^, and syt1^K326,327A^, respectively. We used one-way ANOVA, followed by Dunnett’s test to compare mutants against WT Syt1. *, **, and *** indicate p<0.05, p<0.01, and p<0.001, respectively.

### Polybasic mutations in either C2 domain impairs membrane binding

To uncover the molecular mechanism by which the polybasic mutations impair evoked neurotransmitter release, we measured the membrane binding affinities and kinetics of both mutant C2AB domains, using a recently developed single-molecule approach^48^ based on optical tweezers (OT). A single Syt1 C2AB domain was tethered between a silica bead and a polystyrene bead, forming a dumbbell in solution suspended by optical traps (Figure 5A). The silica bead was coated with a lipid bilayer^48^ that contained 85 mol% POPC, 10 mol% DOPS, 5 mol% PI(4,5)P_2_ and 0.03% mol% biotin-PEG-DSPE to mimic the plasma membrane. The Syt1 C2AB domain was attached to the lipids through biotin-streptavidin interaction via via an N-terminal 73 amino acid flexible polypeptide linker and pulled from the C-terminus via a 2,260 bp DNA handle. Dynamic C2AB-membrane binding and unbinding events were detected by the corresponding changes in the extension of the protein-DNA tether. We previously determined the membrane binding affinities of the wild-type tandem C2AB domain (Figure 5B) and the individual C2B domain of Syt1^48^. Here we applied the single-molecule approach to characterize membrane binding of Syt1 with the polybasic patch mutations in either C2 domain and a calcium-binding mutation in the C2B domain. All our experiments were conducted in the presence of 100 µM Ca^2+^ in the solution.

**Figure 5.**
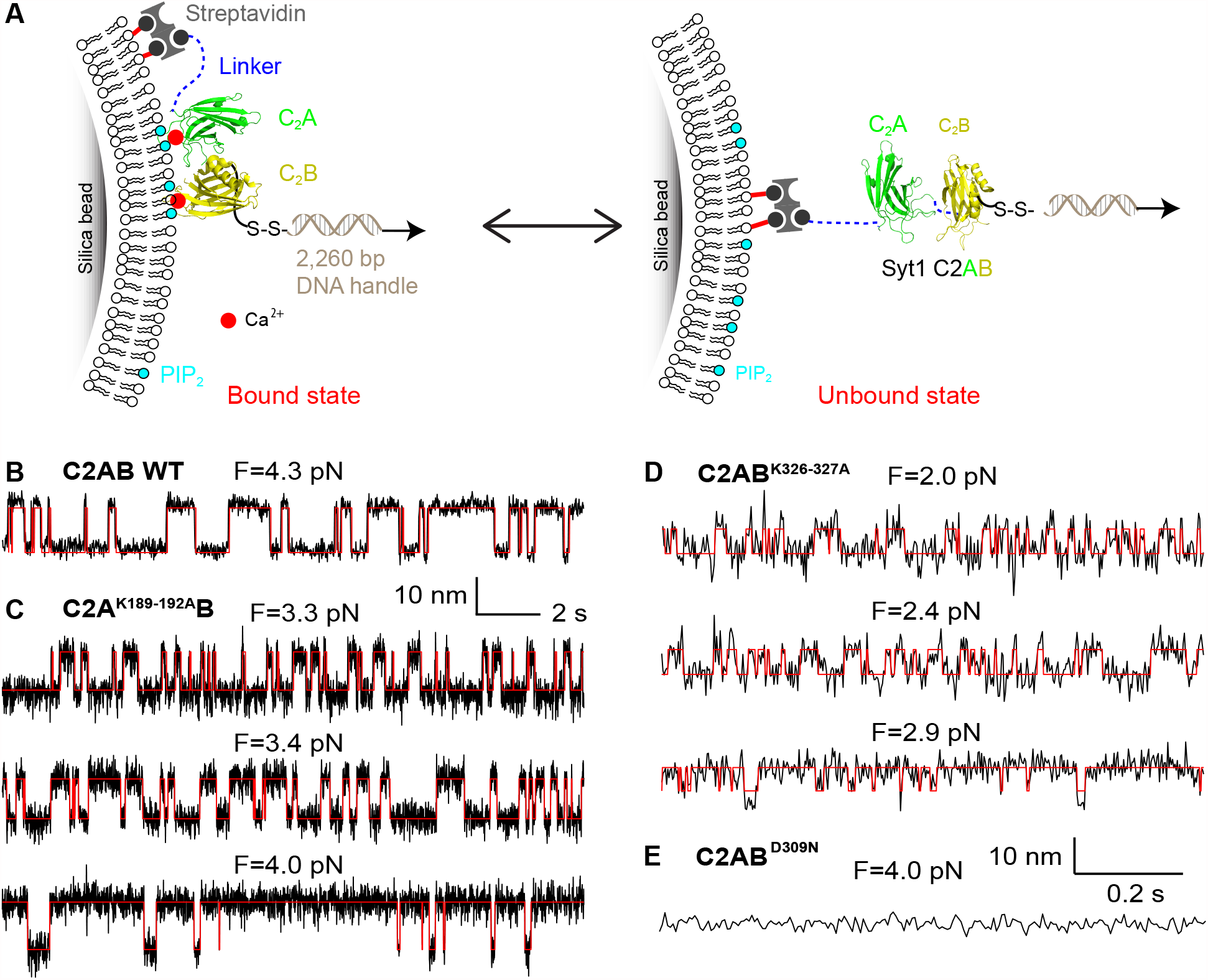
The neutralization mutations in Syt1 C2AB domain impair its membrane binding as revealed by optical tweezers. **A**. Schematic of the experimental setup. The Syt1 construct was directly attached to the supported bilayer *via* biotin-streptavidin interactions at the N-terminus and crosslinked to a DNA handle *via* a disulfide bond. The other end of the DNA was attached to a polystyrene bead (not shown). Membrane binding and unbinding of the C2AB domain was detected by the corresponding extension change of the protein-DNA tether. The bilayer is composed of 85 mol% POPC, 10 mol% DOPS, 5 mol% PI(4,5)P_2_ and 0.03% mol% biotin-PEG-DSPE. **B-E**. Extension-time trajectories at the indicated constant mean forces showing dynamic C2AB binding of wild-type Sty1 C2AB (**B**), C2A^K189-192A^B (**C**), or C2AB^K326-327A^ (**D**), or no membrane binding of C2AB^D309N^ (**E**). Note that the extension-time trajectories in B and C share the same scale bars, so as the data in D and E.

We bacterially expressed and purified the soluble C2AB domains with a polybasic patch mutation in the C2A (C2A^K189-192A^B) or the C2B domain (C2AB^K326-327A^). We detected reversible membrane binding and unbinding transitions of C2A^K189-192A^B in the force range of 2.6-4.2 pN, as is indicated by the two-state extension flickering at constant mean forces (Figure 5C). Here the states with higher and lower average extensions represents the unbound state and the membrane-bound state, respectively. The state probabilities and binding and unbinding rates were determined by hidden-Markov modelling^49^. As expected, the probability of the mutant C2 domain being in the unbound state increases as force increases, while the unbinding rate and binding rate approximately exponentially increases and decreases, respectively (Figure 5C and Figure 6). The analysis also revealed an equilibrium force with equal probabilities being in both states that represents the membrane binding affinity of the protein. C2A^K189-192A^B has a smaller equilibrium force (3.5 pN) than that of the wild-type Syt1 (4.7 pN) (Table 1), which indicates that neutralization of the basic patch in C2A reduces the membrane binding affinity of Syt1. Detailed data analysis^48,49^ revealed that C2A^K189-192A^B binds the membrane with an energy of 9.1 (± 0.9) k_B_T (mean ± SEM), compared with the 12.8 (± 0.8) k_B_T binding energy for the wild-type C2AB. The reduction in the binding energy is mainly caused by a decrease in the membrane association rate constant, with insignificant change in the dissociation rate constant. These observations suggest that the four lysine residues in C2A help Syt1 C2AB domain bind to the membrane through their long-range electrostatic interactions with the membrane, and neutralization of the basic residues reduces the binding rate constant. We similarly measured the membrane binding affinity and kinetics of C2AB^K326-327A^. Neutralization of the two lysine residues in C2B significantly reduces the equilibrium force to 2.1 (± 0.4) pN and membrane binding energy to 5.9 (± 0.4) k_B_T (Figure 5D Figure 6). Interestingly, the mutations decrease the binding energy mainly by increasing the dissociation rate constant, with only a small decrease in the association rate constant (Table 1). Finally, we tested membrane binding by C2AB^D309N^ as a control that impairs Ca^2+^-dependent Sty1 C2AB membrane binding^50^ and neurotransmitter release^20^. Our single-molecule assay did not detect any membrane binding by C2AB^D309N^ (Figure 5E), confirming that calcium promotes membrane binding of the C2AB domain. In conclusion, both the C2A domain K189-192A and the C2B domain K326-327A mutations likely impair neurotransmitter release by reducing membrane binding of the Sty1 C2AB domain required for release.

**Figure 6.**
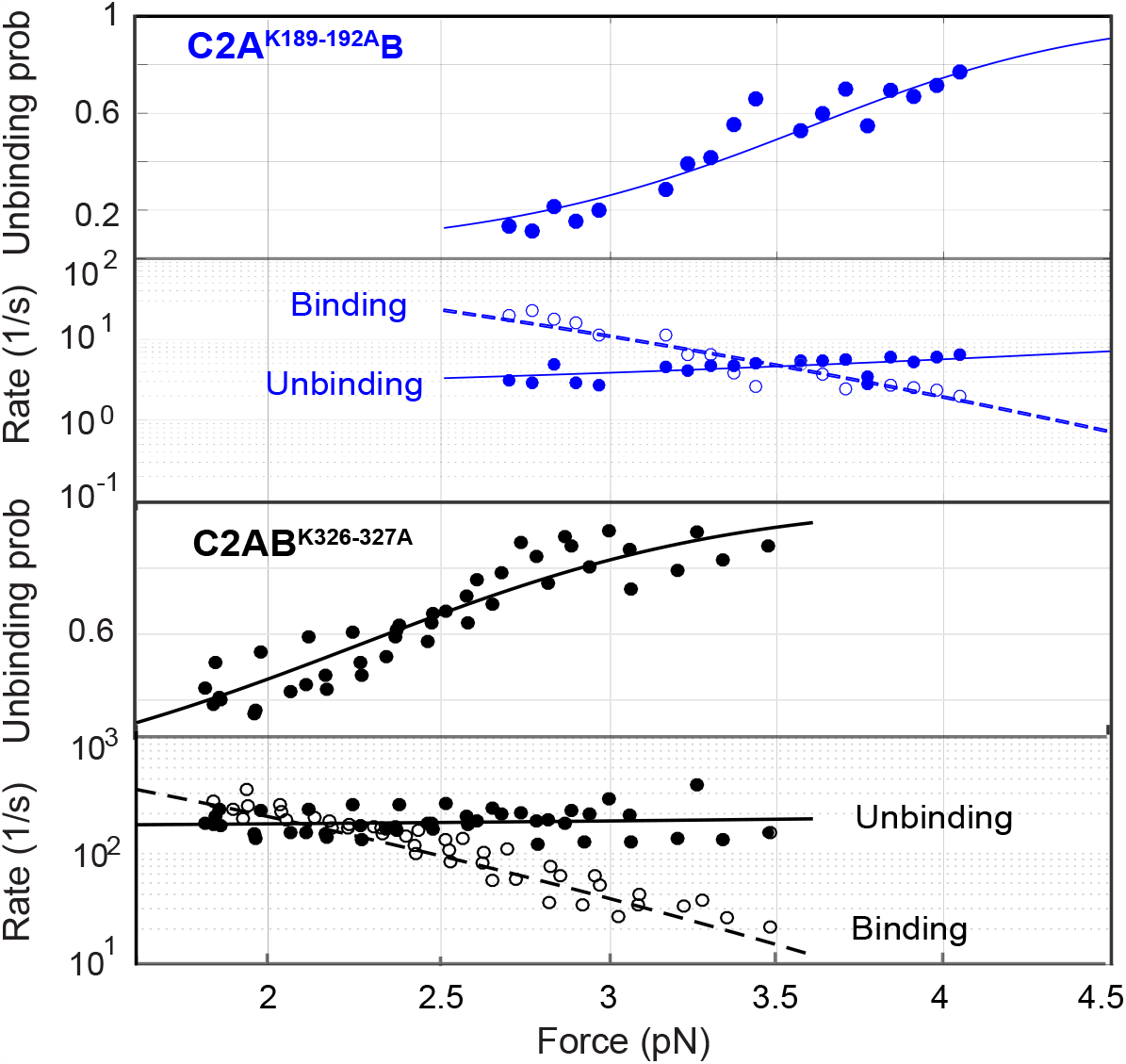
Force-dependent C2AB unbinding probabilities and binding and unbinding rates. All the experimental measurements (symbols) are simultaneously nonlinearly fit by a theoretical model to account for the effect of force on protein binding and unbinding (curves) to derive the membrane binding energy and the rate constants (see Methods) for Syt1 C2A^K189-192A^B (top two rows) and C2AB^K326-327A^ (bottom two rows).

**Table 1.** Comparison of membrane binding energy and kinetics of wild-type and mutant Sty1 C2AB domains. The flexible tether linking the C2 domain(s) to the bilayer on the silica bead increases the likelihood of rebinding, affecting the apparent binding energy. We used a previously developed model to account for this effect, as in Lu et al.^48^.

| Syt1 | Equilibrium force (pN) | Binding energy with tether ($E_b$) ($k_B T$) | Binding energy without tether ($E_{on}$) ($k_B T$) | $\text{Log}_{10}[k_b]$ ($s^{-1}$) | $\text{Log}_{10}[k_{on}]$ ($M^{-1}s^{-1}$) | $\text{Log}_{10}[k_{ub}]$ ($s^{-1}$) |
| --- | --- | --- | --- | --- | --- | --- |
| WT | 4.7 (0.2) | 10.8 (0.8) | 12.8 (0.8) | 4.6 (0.4) | 5.4 (0.4) | 0 (0.2) |
| K189-192A | 3.5 (0.6) | 7.1 (0.9) | 9.1 (0.9) | 3.5 (0.2) | 4.3 (0.2) | 0.4 (0.3) |
| K326-327A | 2.1 (0.4) | 3.9 (0.4) | 5.9 (0.4) | 3.9 (0.1) | 4.7 (0.1) | 2.2 (0.1) |

## DISCUSSION

Two lines of evidence led to the idea that the C2B domain is the functionally more important C2 subunit of Syt1, with the C2A domain mostly playing a faciliatory role. First, impairing calcium binding to C2A has a milder effect than impairing calcium-binding to the C2B domain^3,16-20^.

Second, the C2B domain of Syt1, but not the C2A domain, interacts with the SNARE proteins^22,24,25,27,29^ (but see 23) that drive membrane fusion^51^. However, recent work challenged the merely facilitatory role attributed to C2A, as it was found that the C2A domain is essential for evoked release in flies^35,37,38^ and in mice^39^. Here we asked what feature of the C2A domain may be responsible for its critical role.

Since previous work suggested that calcium binding to the C2A domain may not be an essential functional feature of this C2 domain, we explored the role of another potentially important region, a highly conserved stretch with four tandem lysines (residues 189-192). The C2B domain similarly possesses a conserved polybasic patch on one side (residues 324-327). Separate functional studies of these two polybasic regions lead to inconsistent, or even conflicting results. Neutralization of the C2A polybasic region through mutagenesis only increased spontaneous release in the *Drosophila* larvae NMJ^40^, while injection of peptides into the squid giant nerve terminals^41^ or PC12 cells^42^ suggested the C2A polybasic patch is essential for evoked release. Charge inversion or neutralization mutations of the C2B polybasic region resulted in 40-50% reduction in evoked release but with divergent effects on spontaneous release at the *Drosophila* NMJ^30,31^ and in mouse hippocampal neuronal cultures^32-34^, or a 5-6-fold reduction in evoked release accompanied with a ∼60% reduction in the RRP size, also in mouse hippocampal neuronal cultures^34^. These discrepancies can be due to differences in the species or preparations used, but since there is no consensus among these results, we decided to test the roles of both C2 domain polybasic patches side-by-side to reveal their individual contributions in mammalian synapses.

We found that evoked release was reduced ∼6 and >10-fold in *syt1-/-*mouse cortical neuronal cultures expressing charge-neutralization mutations in the C2A and C2B polybasic patches, respectively. These severe reductions were not due to differences in expression levels or the sizes of the readily releasable pools. Spontaneous release was not significantly affected by the mutations. Thus, both polybasic patches in C2A and C2B are important for evoked release in mammalian synapses. Because the RRP sizes were not different, the most likely explanation for the reduction in evoked release in the mutants was a reduction in release probability.

We reasoned that the effects of the C2A and C2B domain polybasic patch mutations we observed on evoked release could be due to the disruption of putative direct interactions of these patches with acidic membranes. To probe these interactions, we used a single-molecule assay developed recently^48^. Bulk assays that probe protein-membrane interactions are convenient and very valuable but suffer from a number of challenges. First, they often cannot resolve intermediates, energetics, and kinetics of protein-membrane binding due to difficulties in synchronizing the reactions and in applying force to proteins or membranes^52^. Second, Syt1 C2 domains can bridge membranes^53-55^ and/or oligomerize^12,56,57^, likely affecting the bulk measurements. Such interactions may well be physiologically important, but their effects are difficult to disentangle from direct binding-unbinding events in bulk experiments. Our single-molecule measurements avoid possible complications from membrane bridging, since there is only one membrane to bind to. Furthermore, multimerization of Syt1 C2B domains are absent in our assay. Thus, single-molecule measurements directly probe C2 domain-membrane interactions in the absence of potential challenges faced in bulk measurements.

The C2B domain polybasic patch is known to bind acidic lipids, preferably PI(4,5)P_2_, and this interaction can be detected in bulk assays in the absence of calcium^11,14,15^. By analogy, the C2A domain polybasic patch could contribute to membrane binding. Indeed, using single-molecule atomic force microscopy, Takahashi et al.^58^ reported reduced binding of the C2AB domain to supported bilayers when thee C2A polybasic patch was neutralized, though no binding energies could be extracted from the measurements carried under far-from-equilibrium conditions. In addition to the two polybasic patches, Syt1 C2A and C2B domains can also bind membranes through their calcium-binding loops. When bound, calcium ions coordinate between highly conserved aspartates in the C2A and C2B calcium-binding loops and head-groups of negatively charged lipids in the membrane^5^. Hydrophobic residues at the tips of the calcium-binding loops in both C2 domains penetrate into the membrane in the presence of calcium, fortifying membrane binding^5,12,13^. Thus, there are at least four membrane binding sites on Syt1, two on each C2 domain: a calcium-dependent site and a polybasic patch. Our results show that the contributions of these sites to overall binding is not additive, suggesting highly cooperative interactions.

We note that some membrane binding activities that the single-molecule OT assay fails to detect have been reported using ensemble measurements. Notably, membrane-binding of a C2AB construct carrying a mutation that prevents calcium binding to the C2B domain can be detected in bulk assays in the presence of calcium^59^ (but see 50), presumably via calcium-dependent C2A-membrane interactions^5,13,28^. Similarly, membrane binding for the C2AB domain can be detected using bulk liposome binding assays in the absence of calcium^11,14,15^, but not in our OT assay^48^. Finally, in the presence of calcium, the C2A domain avidly binds membranes containing phosphatidylserine (PS) in bulk experiments^5,13,28^, but the OT assay fails to detect the interaction^48^. These apparent discrepancies are actually expected for several reasons. First, bulk assays are often more sensitive to detect weak interactions, because the ensemble average of signals from a large number of molecules are detected^60^. Second, the OT assay measures dynamic binding and unbinding events under a load and these must occur within a certain range of force and time scales to be detectable. By contrast, many bulk measurements probe equilibrium properties. That is, even if a bulk equilibrium assay reports a low dissociation coefficient *K*_*d*_, the dynamics may not be readily detectable in the OT assay. More subtle is the effect of applied load on dissociation kinetics: if unbinding is accelerated under sufficiently low applied forces, binding-unbinding events may not be detectable in the OT assay, even if such events can be detected in single-molecule fluorescence assays where no load is applied. Third, Syt1-membrane interactions may be promoted by high membrane curvature^61,62^, such that binding to small liposomes in bulk experiments may be stronger than binding to a relatively flat supported bilayer in our OT assay. Finally and perhaps most likely, some of these apparent discrepancies may be due to the relatively low amount of PS (10 mole %) we included in the bilayers.

The results above are important for our interpretation of the contribution of the C2A and C2B domain polybasic patches to Syt1-membrane interactions. In particular, although the C2A domain binds acidic membranes (detected using bulk assays), binding has not been reported in the absence of calcium, at least in the presence of moderate amounts of PS^5,13,28^, leading to the idea that the C2A polybasic patch does not contribute significantly to the overall Syt1-membrane interactions. Our results clearly demonstrate that both the C2A and the C2B domain polybasic patches significantly enhance calcium-dependent lipid binding of the C2AB domain. Given the preference of the C2A domain for binding PS over PI(4,5)P_2_, and the fact that the inner leaflet of the plasma membrane contains ∼20 mole% PS^63^, about twice the amount we used in our OT assay, it is likely that the C2A domain polybasic patch-plasma membrane interactions are even stronger *in vivo* and physiologically relevant.

How can we explain the cooperativity between the four binding sites? The simplest idea is that multiple attractive interactions increase the binding rate and dramatically slow the overall kinetics of unbinding, even if individual binding sites each contribute a relatively small interaction energy. This is a well-known phenomenon for polymer adsorption to surfaces^64^. Even if individual segments on a random polymer adsorb onto a surface with a small binding energy of order k_B_T and come off rapidly due to thermal energy, as one segment unbinds, another can bind, maintaining multiple binding sites occupied at any given moment. For long polymers the probability that all segments unbind at the same time is very small and adsorption is essentially irreversible. In the case of Syt1 C2AB domains, the cooperativity of the binding sites is certainly more complicated, as the domains are well folded and the surface they bind is soft, allowing multiple bound-state configurations. This complexity is likely reflected in our finding that neutralizations of the C2A or the C2B polybasic sites both lead to a reduction of the overall binding energy, but through different pathways. In the former case the binding rate was >10-fold slower compared to wild-type C2AB, with a modest effect on the unbinding rate. For the latter, the unbinding rate was accelerated two orders of magnitude, with a smaller effect on the binding rate.

Our results are broadly consistent with the observation that Syt1 mutations that inhibit calcium-binding to the C2A domain while partially mimicking the charge-neutralization by calcium-binding (D232N) do not reduce synchronous release, and even enhance it^19,65,66^. In these mutants, the calcium-dependent binding site on C2A would be replaced by a partial calcium mimic, turning it to a calcium-independent binding site (note however, that these mutants do not display increased calcium-independent binding to anionic lipids^12^, so the actual picture is likely to be more complex). By contrast, inhibiting calcium-binding by an aspartate-to-glutamate (D→E) substitution removes the calcium-dependent binding site on C2A altogether, and results in a dramatic decrease in evoked release^35^. The fact that similar D→N substitutions in the C2B domain calcium-binding site result in severe inhibition of evoked release^17^ likely reflect the fact that the C2B domain binds the SNARE complex and that calcium-binding to the C2B domain likely leads to a reorientation of the C2B-SNARE complex^45,67,68^, its dissociation^69,70^, or some other specific re-arrangement required for triggering membrane fusion.

Cooperativity between the C2 domains of Syt1 was noted long ago, but the nature of this cooperativity has proven difficult to pin down^5,71^. Our results provide new insights into this question and show that both inter- and intra-C2 domain binding sites of Syt1 contribute to membrane binding in a highly cooperative manner.

## MATERIALS AND METHODS

### Neuron preparation, syt1 lentivirus production and transduction

Primary cortical neurons from heterozygous syt KO breeders were isolated at postnatal day 0-1. All procedures are in accordance with the guidelines of NIH for the Care and Use of Laboratory Animals. The protocols were reviewed and approved by the Animal Care and Use Committee (ACUC) at the University of Wisconsin-Madison. In brief, cortices were dissected from mouse brain and digested for 25 min at 37 °C in 0.25% trypsin-EDTA (Corning). After mechanical dissociation, cortical neurons were plated on 12 mm glass coverslips, which surfaces were pretreated with poly-D-lysine (Thermofisher) for at least one hours at RT. Cultures were maintained in Neurobasal A (GIBCO) based medium with B27 (2%, Thermofisher) and GlutaMAX (2 mM, GIBCO) at 37 °C incubator.

For virus infection experiments, WT and mutant forms of syt1 DNA were subcloned into a FUGW transfer plasmid (Addgene plasmid # 14883) modified with a synapsin promoter and an IRES-expressed soluble GFP marker. Lentiviral particles were generated as previously described^39^. In brief, Transfer plasmids which carry WT and corresponding mutated syt1 were co-transfected with packaging and helper (pCD/NL-BH*ΔΔΔ and VSV-G encoding pLTR-G) plasmids into HEK 293/T cells. Lentivirus was collected from the media 48–72 h after transfection and concentrated by ultracentrifugation. These viruses were first titrated and then infected neurons on day-in-vitro (DIV) 3-5.

### Electrophysiology

Cortical neurons (DIV 14-19) with GFP fluorescence were patched at room temperature using a Multiclamp 700B amplifier (Molecular Devices). Recording pipettes were pulled from borosilicate glass (Sutter Instruments) and filled with the pipette internal solution composed of (in mM): 130 KCl, 1 EGTA, 10 HEPES, 2 ATP, 0.3 GTP, and 5 sodium phosphocreatine, Ph 7.35 and 290 mOsm, which generate a 3–5 MΩ resistance in a bath solution containing (in mM): 128 NaCl, 5 KCl, 2 CaCl2, 1 MgCl2, 30 D-glucose, and 25 HEPES, pH 7.3 and 305 mOsm. Patched neurons for data collection were held at −70 mV with access resistance (*R*_*a*_) less than 15 MΩ at any time point. GABAA receptor mediated currents were pharmacologically isolated by including D-AP5 (50 µM, Abcam) and CNQX (20 µM, Abcam) in the bath solution. Data were obtained using a Digidata 1440A (Molecular Devices) and Clampex 10 software (Molecular Devices) at 5 kHz. IPSCs were captured with 5 mM QX-314 in pipette solution and evoked by a single stimulus via a concentric bipolar electrode (FHC, 125/50 µm extended tip), which were places ∼300 µm away from the neural soma. The stimulating amplitudes (∼ 0.5 to 1 mA) and position of stimulator tip were adjusted per recording to acquire the stable IPSC currents. For miniature IPSC measurements, tetrodotoxin (TTX, 1 µM) was included in the bath solution to inhibit all action potentials. 300 seconds of data were recorded for each cell and miniature events were identified and analyzed by the template matching algorithm in Clampfit.

The RRP (readily releasable pool of vesicles) size was quantified by osmotic shock. In brief, 0.5 M sucrose in bath solution were puffed by a Picospritzer III (Parker Hannifin) through a fused silica needle (28 gauge, WPI), which placed ∼500 µm away from the soma of aimed neuron. Recorded neurons were fully covered by sucrose and treated for 15 s, yielding a distinct fast and slow phase of release. To avoid tonic effect, only the fast components were utilized to determine RRP size.

### Western blotting

Cortical neurons (14-19 DIV) expressing WT or mutant forms of syt1 were treated with lysis buffer (50 mM Tris pH 8.0, 150 mM NaCl, 2% SDS, 0.1% Triton X-100, 10 mM EDTA) supplemented with protease inhibitors (complete mini EDTA-free, Roche, 1 tablet / 50 mL lysis buffer). The lysed neurons were mixed with 4X Laemmli sample buffer and heated at 70°C for 10 minutes (stored at −20°C until use). The total proteins were separated by SDS-PAGE gel, and then transferred to PVDF membrane. The membranes were blocked with 5% milk and incubated with an anti-syt1 primary antibody (3 µg / ml; Developmental Studies Hybridoma Bank; Cat#: mAb 48) and a GAPDH primary antibody (1:1000; Cell signal; Catalog # 2118) in TBS-T with 1% milk at 4°C overnight. After washing in TBS-T, blots were incubated with Goat anti-Mouse IgG2b Cross-Adsorbed Secondary Antibody, HRP (1:2000; Invitrogen; Catalog # M32407) or Goat Anti-Rabbit IgG (H L)-HRP Conjugate (1:1000; Bio-Rad; Catalog # 172-1019) in TBS-T for 1 hr at room temperature, washed in TBS-T, and then imaged with a CCD gel imaging device (GE).

### Immunocytochemistry and confocal microscopy

Cortical neurons (DIV 14-18) on coverslips were fixed by incubation with pre-warmed 4% paraformaldehyde for 10 minutes at 37°C. Cell membranes were permeabilized by 0.2% saponin for 10 minutes, rinsed in PBS, and blocked for 1 hour at room temperature in PBS buffer supplemented with 5% normal goat serum, 5% BSA, and 0.02% saponin. Cells were then incubated with syt1 monoclonal antibody (3 µg / ml; Developmental Studies Hybridoma Bank; Cat#: mAb 48) and synaptophysin 1 primary antibody (1:500; guinea pig pAb; Synaptic Systems; Catalog # 101 004) in PBS containing 1% BSA and 0.02% saponin at 4°C overnight. Cells were washed in PBS containing 0.02% saponin and incubated with Goat anti-Mouse IgG2b Cross-Adsorbed Secondary Antibody, Alexa Fluor 594 (Invitrogen; A-21145; 5 µg / ml) and Goat anti-Guinea Pig IgG (H+L) Highly Cross-Adsorbed Secondary Antibody, Alexa Fluor 647 (Invitrogen; Catalog # A-21450; 5 µg / ml) for 1 hour at room temperature. Stained neurons were imaged on a Zeiss LSM 880 confocal microscope equipped with a 60x oil objective. Syt 1 and synaptophysin colocalizations were quantified by Pearson’s correlation coefficients using the Coloc 2 plugin in FIJI^72^. Eight or nine slides from at least 2 separate breeders at each group were imaged.

### Protein expression and purification

The wild-type rat Syt1 construct used for the single-molecule assay and its preparation were described in detail previously^48^. The Syt1 mutants were created from the wild-type Sty1 construct using PCR mutagenesis and prepared similarly to wild-type Syt1. Briefly, all the Syt1 constructs contain an N-terminal Avi-Tag, a flexible polypeptide linker, the C2AB domain, and a unique cysteine residue added to the C-terminus. The DNA coding sequences of Syt1 proteins were cloned into pET SUMO vector which contains a His tag and a sumo protein at the N-terminus. All Syt1 proteins were expressed in BL21 *E. coli* cells and purified using the His-tag. Protein biotinylation was carried *in vitro* using BirA ligase which catalyzes biotinylation of a lysine residue in the Avi-Tag. The SUMO tag was then removed by the SUMO protease. The purified Syt1 protein was crosslinked to the DNA handle by mixing the two at a 50:1 molar ratio under an oxidization condition.

### Single-molecule assay for Syt1-membrane interactions

The dual-trap high-resolution optical tweezers are home-built and calibrated as previously described^73^ Briefly, a 1064 nm laser (CW, Nd: YVO4 laser with a maximum power of 5 W) beam is expanded by ∼10 fold in diameter by two telescopes. Between the telescopes, the laser beam is split into two orthogonally polarized beams. One beam is steered by a mirror attached to a nano-positioning stage, which is used to move the optical trap in the sample plane. The two beams are then focused by a water immersion 60× objective (NA = 1.2) to form two optical traps inside the central channel of a microfluidic flow cell. The outgoing laser beams are collected and collimated by a second identical objective, split again by polarization, and projected onto two position-sensitive detectors to detect bead positions using back-focal-plane interferometry^73^.

## ACKNOWLEDGEMENTS

We thank the members of the Karatekin, Chapman, and Zhang labs for valuable discussions. We are grateful to D. Zenisek (Yale University) for discussions and feedback on the manuscript. We acknowledge funding from the National Institutes of Health, National Institute of Neurological Disorders and Stroke (grant R01 NS113236 to EK and R35 NS097362 to ERC), National Institute of General Medical Sciences (R35 GM131714 to YZ), National Eye Institute (R01EY010542 to EK), and National Institute Of Mental Health (R01 MH61876 to E.R.C.). E.R.C. is an Investigator of the Howard Hughes Medical Institute. The funders had no influence in the design, execution, and interpretation of the study.

## AUTHOR CONTRIBUTIONS

ZW, LM, and EK conceived the study. ZW, NAC, and JZ performed the experiments with cortical neuronal cultures. LM performed all single-molecule experiments. ZW, LM, NAC, JZ, YZ and EK analyzed data. EK, YZ, and ERC acquired funding and supervised the work.

